# Similarities in the behaviour of dance followers among honey bee species suggest a conserved mechanism of dance communication

**DOI:** 10.1101/2020.02.02.931345

**Authors:** Ebi Antony George, Smruti Pimplikar, Neethu Thulasi, Axel Brockmann

## Abstract

Group living organisms rely on intra-group communication to adjust individual and collective behavioural decisions. Complex communication systems are predominantly multimodal and combine modulatory and information bearing signals. The honey bee waggle dance, one of the most elaborate forms of communication in invertebrates, stimulates nestmates to search for food and communicates symbolic information about the location of the food source. Previous studies on the dance behaviour in diverse honey bee species demonstrated distinct differences in the combination of visual, auditory, olfactory, and tactile signals produced by the dancer. We now studied the behaviour of the receivers of the dance signals, the dance followers, to explore the significance of the different signals in the communication process. In particular, we ask whether there are differences in the behaviour of dance followers between the 3 major Asian honey bee species, *A. florea, A. dorsata and A. cerana*, and whether these might correlate with the differences in the signals produced by the dancing foragers. Our comparison demonstrates that the behaviour of the dance followers is highly conserved across all 3 species despite the differences in the dance signals. The highest number of followers was present lateral to the dancer throughout the waggle run, and the mean body orientation of the dance followers with respect to the waggle dancer was close to 90° throughout the run for all 3 species. These findings suggest that dance communication might be more conserved than implied by the differences in the signals produced by the dancer. Along with studies in *A. mellifera*, our results indicate that all honey bee species rely on tactile contacts between the dancer and follower to communicate spatial information. The cues and signals that differ between the species may be involved in attracting the followers towards the dancer in the different nest environments.

## Introduction

Organisms have evolved various ways to communicate amongst themselves [1]. Communication involves indirect cues and direct signals and varies in its complexity [2,3]. Complexity in communication systems correlates with social group complexity [4, but see 5]. Social communication mechanisms consist of multiple signal channels which can be of the same modality, e.g. ant pheromone trails, or different modalities, as in the case of ritualised courtship signals in birds [6,7]. In these communication systems, the different signals are either equally informative, or one of the signals contains the information and the others act as modulators, enhancing the effect or spread of the signal [8]. Finally, environmental factors and plasticity in the signal can lead to divergence in signals across closely related species [9–11].

One of the most elaborate types of social communication in invertebrates is the honey bee waggle dance used by foragers returning from profitable food sources to recruit nestmates [12]. The waggle dance motivates foragers to fly out and in addition encodes spatial information about the food source [13,14]. Each waggle dance consists of multiple circuits and one circuit contains two phases; a straight walking phase in which the dancer waggles its abdomen back and forth (the waggle run or the waggle phase) and a circular walking phase which brings the dancer back towards the point of origin of the first phase (the return phase). The duration of the waggle run corresponds to the distance to the food source [12,14]. In *Apis mellifera*, in which foragers dance in the dark on vertical combs, the body orientation of the dancer with respect to the vertical (gravity) axis during the waggle run corresponds to the direction of the food source from the hive with respect to the sun’s azimuth [12]. Further, the duration of the return phase corresponds to the reward value of the food source as perceived by the forager [15]. For food sources very close to the hive, the dance circuit becomes nearly circular with a very short waggle run [16].

Cues and signals from the environment and nestmates can modulate the probability and intensity of dance behaviour [15,17–19]. Interactions with nestmates in the hive inform nectar foragers about the colony food stores and the nectar influx into the colony [18,20,21]. In addition, interactions with other foragers provide information about predation and overcrowding at the food source [22,23]. An individual forager’s dance activity is modulated by the perceived reward value of the food source along with information from these interactions [19]. This in turn drives recruitment to each food source proportional to its relative reward value, which leads to an efficient distribution of the colony’s foraging force [24]. Thus, the waggle dance acts as the primary regulatory mechanism of the colony’s recruitment activity in addition to its role in the efficient spatial distribution of the colony’s foragers.

Although extensive research has been done on the honey bee waggle dance behaviour, the mechanism underlying the transfer of spatial information has remained elusive [25,26]. The experimental difficulty lies in determining which of the multiple dances followed is used to obtain information [24] and in tracking whether the follower visited the indicated food source. Moreover, recent studies showed that followers can choose to rely on either the information from the dancer or their own memory [27,28]. Currently, there are two major hypotheses on which signals dance followers use to obtain spatial information from the dancer. The “tactile hypothesis” proposes that dance followers use tactile signals, associated with physical contact between the dancer and follower [29,30] or even mechanosensory signals, associated with the air flow caused by a dancer’s vibrating wings [31,32]. The “follow hypothesis” suggests that followers obtain information from dancers by following the path of the dancer from behind [33]. In this case, followers receive the dance information from their own body positions and walking paths to calculate the direction and distance of the food source being advertised. Studies on the mechanism of spatial information transfer in *A. mellifera* offered preliminary evidence for both hypotheses [29,33]. Instead, comparative studies on dance communication including the behaviour of the dance followers in different honey bee species might help to decide the controversy [34].

Interestingly, there are characteristic differences between the species in the combination of signals generated by the dancer [34–39]. Dances of the dwarf honey bees (e.g. *A. florea*), which take place on a horizontal surface exposed to the sun, include a conspicuous visual signal (raised abdomens) but no auditory signals [40]. The giant honey bees (e.g. *A. laboriosa* and *A. dorsata*) usually dance on the vertical surface of the bee curtain exposed to the sun [38]. Dances of the diurnal *A. laboriosa* are silent [41], whereas those of the cathemeral *A. dorsata* contain auditory signals [42]. All cavity nesting species investigated (e.g. *A. mellifera*, *A. cerana* and *A. nigrocincta*), which perform dances in the hive, produce auditory signals [38]. However, it is unclear whether these differences in dance signals imply the evolution of different mechanisms for information transfer across the genus or whether the mechanism is the same with the different signals and cues serving to attract followers [43,44].

We performed a comparative study to explore the dance follower behaviour of Asian honey bees and to test if possible differences in dance follower behaviour might correlate with differences in the signals and cues produced by the dancer. Parallel changes in the behaviour of the dancer and dance follower would be a strong argument for the evolution of different mechanisms of information transfer across the genus *Apis*. On the other hand, a high degree of similarity in the behaviour of the dance follower would suggest that the major mechanism of information transfer is conserved. Further, we monitored the spatial positions of the followers to obtain evidence for either of the two proposed hypotheses regarding the mechanism of information transfer. If followers arrange themselves towards the side of the dancer, then it is likely that they use tactile cues (“tactile hypothesis”) to obtain information about the food source [29]. In contrast, if followers orient themselves behind the dancer, they are more likely using their own body orientation (“follow hypothesis”) to determine the spatial position of the food source [33].

## Methods

### Experimental Location, Colonies and Distance Training

This study is based on analysis performed on videos of waggle dances *A. florea*, *A. dorsata* and *A. cerana* that were recorded as part of another study [45].

The experiments were performed in the Botanical Garden at the University of Agricultural Sciences, Gandhi Krishi Vignana Kendra, Bengaluru (latitude: 13.07, longitude: 77.57). The garden provides dense vegetation cover and hence good optic flow for the foragers (fig S1). Experiments were done with two *A. florea* and *A. cerana* colonies each and one *A. dorsata* colony. The first set of colonies from all 3 species was observed in January - March 2017, and the second *A. cerana* and *A. florea* colony was observed in February - April 2018. For further details on colony preparation, see Kohl *et al.*, 2020 [45].

A brief description of the distance training protocol employed for all three species in Kohl *et al.*, 2020 [45] is provided here. The foragers in all three species were trained along a 500 m transect using an artificial feeder filled with sucrose solution scented with star anise (*Illicium verum*) extract. The sucrose concentration at the feeder was adjusted between 1 and 2.5 M depending on the number of foragers visiting our food source. Individual foragers were paint marked using Uni POSCA Paint markers (Uni Mitsubishi Pencil, UK). The dance activity of the marked foragers was recorded at 1080p and 50 frames per second for one hour each at 100 m, 200 m, 300 m, 400 m and 500 m using a Sony HDR CX260V Handycam (Sony Corporation, Tokyo). In the case of *A. dorsata*, the dance activity was recorded at distances of 100 m, 200 m, 300 m and 400 m, but not at 500 m since foragers did not come to the feeder at this distance.

### Video Analysis

For the video analysis, individual foragers were first shortlisted based on whether they were active at the feeder at multiple distances. We then analysed each dance circuit in the dances by these individuals to determine the duration of the waggle run. The videos were observed frame-by-frame in Virtual Dub 1.10.4 (http://www.virtualdub.org). The first frame in which a focal bee clearly moved its abdomen laterally or dorsoventrally was defined as the start of the waggle run in that circuit. The frame in which the bee stopped waggling its abdomen and started turning to the left or right was defined as the end of the waggle run. The time between the start and the end frames was calculated as the duration of the waggle run.

### Follower Behaviour

We defined dance followers as those bees which positioned themselves within one bee length of the dancer and excluded others who were beyond this distance threshold and not following the dancer [30]. In each run, we focussed on 3 phases (time-points); the Start, Middle and End, based on the waggle run duration we had calculated (fig 1 *a*). This was done to look at whether there was a change in the number of followers as the run progressed [29]. At each time-point, the number of followers present in the following three zones (fig 1 *b*) around the dancer was counted; the anterior zone around the head region of the dancer, the lateral zone near the thorax and abdomen, and the posterior zone behind the abdomen [29].

**Figure 1.**
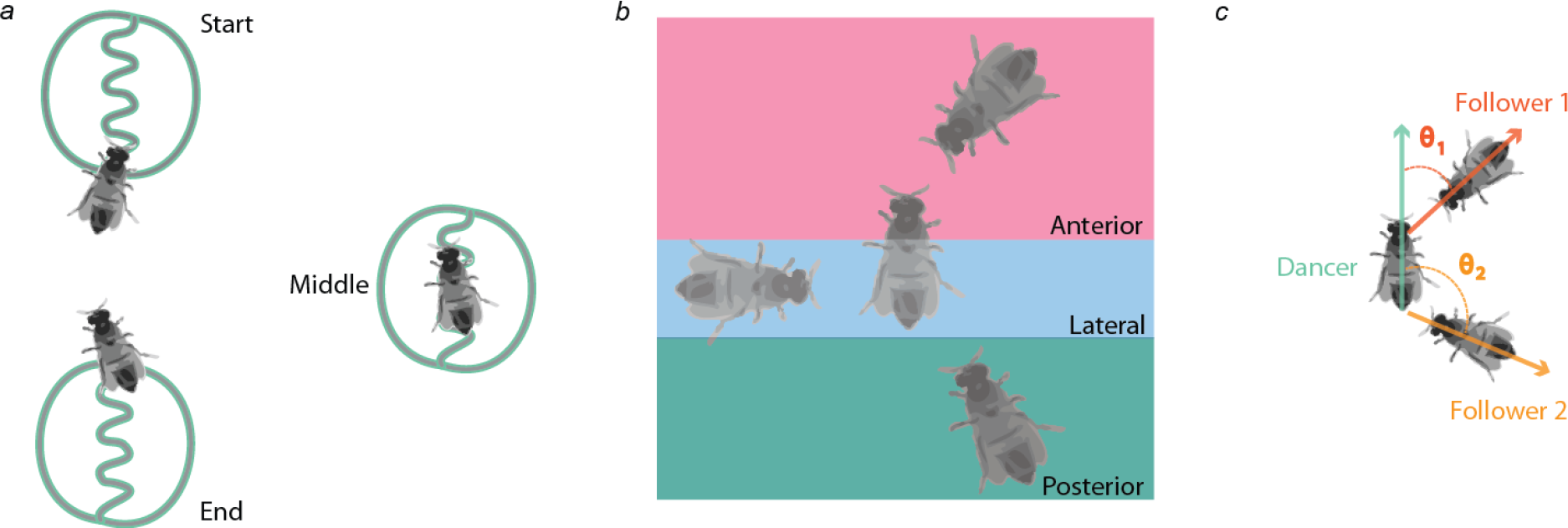
Schematic of the analysis done. (*a*) Each waggle run was divided into 3 phases: the start, middle and end. (*b*) Followers around the dancer were grouped into 3 zones (Anterior, Lateral and Posterior) based on the position they occupied around the dancer (area of the zones in the figure are representative). (*c*) The orientations of the followers with respect to the dancer (θ) were then quantified.

The body angles of the follower with respect to the angle of the dancer were quantified manually using OnScreenProtractor v0.3 (http://osprotractor.sourceforge.net). We first made a dancer vector, pointing in the abdomen to head direction of the dancer. We then made follower vector, pointing from the follower’s head to its abdomen. Finally, the body angle of the follower was quantified as the angle subtended by the follower vector with respect to the dancer vector in the clockwise direction (fig 1 *c*). In total, we calculated 5036 follower positions and body angles (from 330 waggle runs of 41 waggle dances) in *A. florea*, 1363 (from 119 waggle runs of 7 waggle dances) in *A. dorsata* and 4989 (from 411 waggle runs of 35 waggle dances) in *A. cerana* (table S1). Since followers were not individually identifiable, it is possible that some of the followers we counted were the same across multiple runs and dances.

### Statistical Analysis

#### Number of Followers

The dataset of the number of followers was zero inflated (21.17 % zero values) and hence we fit zero-inflated Poisson models [46]. Models were built with different combinations of 4 predictors for the conditional part of the model (table S2): i) zone of dance follower (a categorical variable of 3 levels; Anterior, Lateral and Posterior), ii) phase of waggle run (a categorical variable of 3 levels; Start, Middle and End), iii) species (a categorical variable of 3 levels; *A. florea*, *A. cerana* and *A. dorsata*) and iv) distance (a continuous variable which was scaled with a mean of zero and a standard deviation of 1). For the zero-inflated part of the model, we fit the 3 categorical variables in all models except 5 (due to model convergence errors, see table S2). We then compared the models based on their AIC values and shortlisted those within a cut-off of 0.95 cumulative Akaike weights [47]. Further, we performed multiple comparisons (with Tukey corrections) of the estimated mean number of followers. We focussed on three comparisons: i) between the 3 species, ii) between the different zones within each waggle run phase and iii) between the same zones across the waggle run phases. We did these specific comparisons based on the important predictors in the shortlisted model (table S2 and S3).

#### Orientation of followers

We used circular statistics to analyse the orientation of the dance followers. We first constrained the body angles of all the dance followers to lie between 0° and 180°, by converting all the angles greater than 180° to their mirror images in the 0° - 180° range (e.g., 210° was converted to 150°, 270° to 90° etc). We based this on the assumption that occupying either the left or the right side of the dancer provided similar access to information for the follower. In addition, this prevented us from obtaining biased estimates of the circular mean and length due to potential bimodality in the circular distributions. After checking for unimodality using the Rayleigh test for unimodal departures from uniformity [48], and reflective symmetry [49], we used Fishers Circular Nonparametric test to compare the median angles of the circular distributions [50,51]. We compared the circular distributions for each of the 3 pairs of the waggle run phase to determine which were different from each other.

All the models and the plots were made in R [52] using the RStudio IDE [53]. The GLMMs were fit using the glmmTMB package [54], model selection and averaging were done using the MuMIn package [55] and the model assumptions were checked using the DHARMa package [56]. Multiple comparisons were done using the emmeans package [57]. Circular statistics were done using the circular package [58] and code found in Pewsey et al. 2013.

## Results

### Number of Followers

We found only one model at the 0.95 cut-off level for cumulative sum of Akaike weights based on our model comparisons (table S3). In this model, the important predictors were the species and an interaction between the waggle run phase and the zone of the follower, but not the distance (table S4). In the zero-inflated part of this model, none of the predictors significantly correlated with the number of absences (i.e., in the number of observations where there were no followers), and these results are provided in the supplementary information (table S4).

#### Effect of species

The species had an effect on the number of followers in the conditional model, but there was no interaction between species and any of the other predictors (table S4 and fig 2 *a* and S2). There were fewer followers per run in *A. cerana* as compared to both *A. florea* and *A. dorsata* (estimated mean – *A. cerana*: 1.188; *A. florea*: 1.512; *A. dorsata*: 1.549; t ratio – *A. cerana* vs *A. florea*: 10.705, *p* < 0.001; *A. cerana* vs *A. dorsata*: 7.082, *p* < 0.001). The number of followers in *A. florea* and *A. dorsata* were not significantly different (*A. florea* vs *A. dorsata*: t ratio = −0.621, *p* = 0.809).

**Figure 2.**
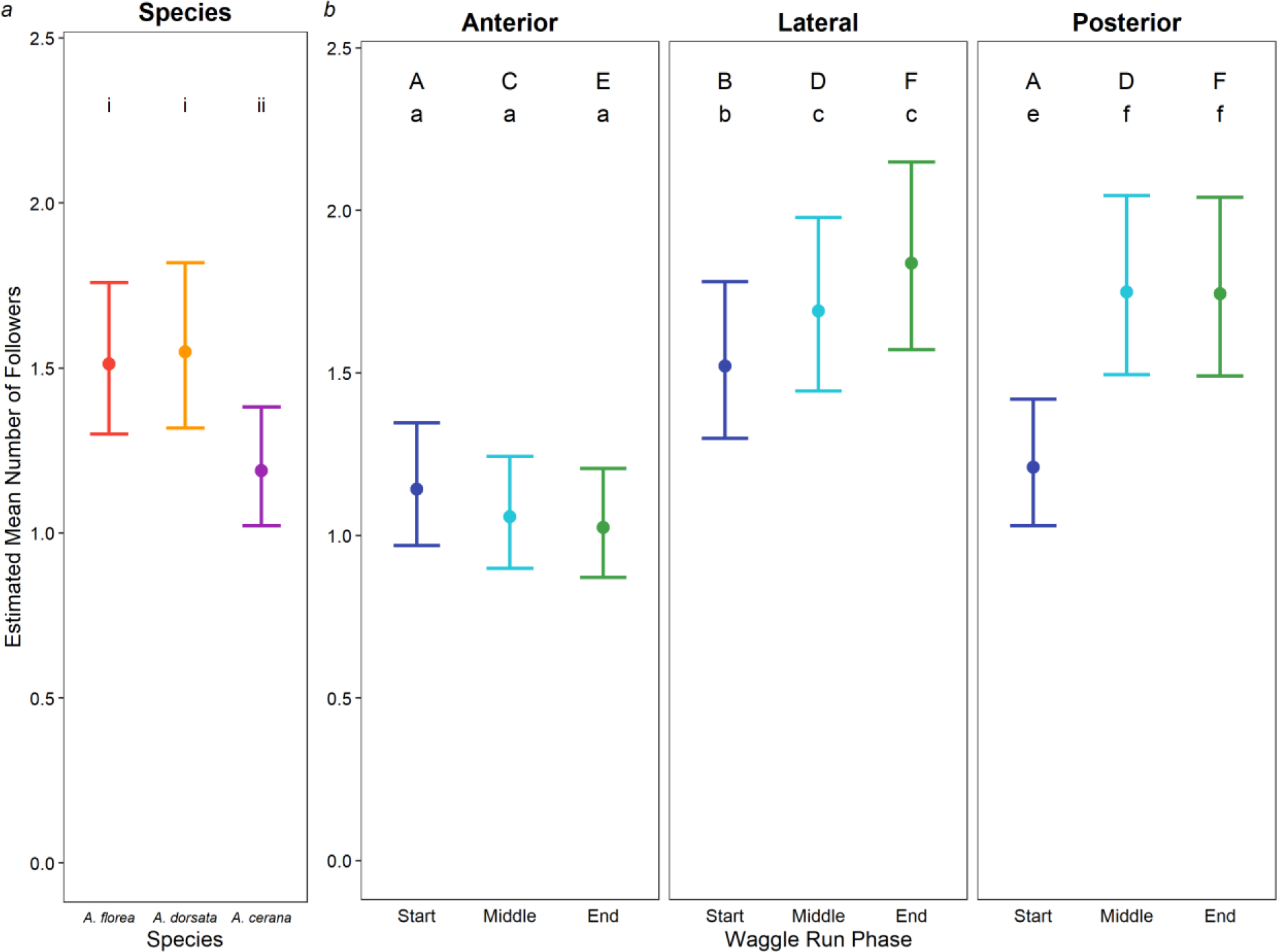
(*a*) The estimated mean number of followers present in the waggle dance of the 3 different species (*A. florea* – red, *A. dorsata* – orange, *A. cerana* – purple; error bars represent 95% confidence intervals). Different roman numerals above the estimates represent significant differences at the *p* < 0.05 level. (*b*) The estimated mean number of followers present in the different zones around the dancer and the different phases of the waggle run (colours based on the waggle run phase: Start – blue, Middle – cyan, End – green; error bars represent 95% confidence intervals). The alphabets above each circle represents results from the multiple comparisons done (estimates with different alphabets were significantly different from each other at the *p* < 0.05 level). Upper case alphabets represent differences in the number of followers present in the same waggle run phase across zones. Lower case alphabets represent differences in the number of followers present in the same zone across different waggle run phases.

#### Effect of waggle run phase and zone of the followers

The waggle run phase and the zone around the dancer had an interactive effect on the number of followers in the conditional model and hence their main effects are not considered (table S4 and fig 2 *b* and S2). Within each waggle run phase, there were differences in the number of followers in the different zones (table 1). At the Start of the waggle run, the number of followers in the Anterior and Posterior zone were similar and significantly lesser than the number of followers in the Lateral zone. In the Middle and at the End of the waggle run, the number of followers in the Lateral and Posterior zone were similar and significantly higher than the number of followers in the Anterior zone.

**Table 1.** Results of the multiple comparisons of the mean number of dance followers in the different zones in each waggle run phase and across waggle run phase in the same zone. The means estimated from the model fitting, as well as the t-ratio and associated *p* value are provided, with comparisons highlighted in italics showing significant differences at the *p* < 0.05 level.

| Predictor | Level | Comparison | Estimated Mean | t ratio | p value |
| --- | --- | --- | --- | --- | --- |
| Waggle Run Phase | Start | *Anterior vs Lateral* | *1.142 vs 1.521* | -6.360 | < 0.001 |
|  |  | Anterior vs Posterior | 1.142 vs 1.209 | -1.213 | 0.446 |
|  |  | *Lateral vs Posterior* | *1.521 vs 1.209* | *5.578* | < 0.001 |
|  | Middle | *Anterior vs Lateral* | *1.057 vs 1.690* | *-11.226* | < 0.001 |
|  |  | *Anterior vs Posterior* | *1.057 vs 1.748* | *-12.110* | < 0.001 |
|  |  | Lateral vs Posterior | 1.690 vs 1.748 | -0.927 | 0.623 |
|  | End | *Anterior vs Lateral* | *1.025 vs 1.838* | *-14.046* | < 0.001 |
|  |  | *Anterior vs Posterior* | *1.025 vs 1.743* | *-12.656* | < 0.001 |
|  |  | Lateral vs Posterior | 1.838 vs 1.743 | 1.478 | 0.301 |
| Zone of Follower | Anterior | Start vs Middle | 1.142 vs 1.057 | 1.597 | 0.247 |
|  |  | Start vs End | 1.142 vs 1.025 | 2.212 | 0.069 |
|  |  | Middle vs End | 1.057 vs 1.025 | 0.654 | 0.790 |
|  | Lateral | *Start vs Middle* | *1.521 vs 1.690* | -2.802 | < 0.001 |
|  |  | *Start vs End* | *1.521 vs 1.838* | -5.125 | < 0.001 |
|  |  | Middle vs End | 1.690 vs 1.838 | -2.333 | 0.051 |
|  | Posterior | *Start vs Middle* | *1.209 vs 1.748* | -9.24 | < 0.001 |
|  |  | *Start vs End* | *1.209 vs 1.743* | -9.169 | < 0.001 |
|  |  | Middle vs End | 1.748 vs 1.743 | 0.072 | 0.997 |

There were differences between each zone in the number of followers across the waggle run phase (table 1). In the Anterior zone, the number of followers were similar across all 3 phases of the waggle run. In the Lateral zone, the number of followers increased as the waggle run progressed, although the number of followers was not significantly different in the Middle and the End of the waggle run (*p* = 0.051). In the Posterior zone, the number of followers increased from the Start to the Middle of the run but did not increase further from the Middle to the End of the run.

#### Effect of distance

Distance had no effect on the number of followers (fig S3). Distance was not a predictor that was present in the short-listed models (table S4). Thus, the main effects of the waggle run phase and the interactive effects of zone and species were similar across all distances.

### Orientation of followers

Even though all the median orientations were close to 90° (see table S5 for full circular summary statistics), the species differed in the change of the median circular orientation of the dance followers from the start to the end of the waggle run (fig 3 and S4). In *A. florea*, the medians of the 3 circular distributions (associated with the Start, Middle and End of the waggle run) were significantly different from each other (median – Start: 86.69; Middle: 108.43; End: 100.33; Fisher test statistic – Start vs Middle: 58.898, *p* < 0.001; Start vs End: 25.878, *p* < 0.001; Middle vs End: 10.109, *p* = 0.001). In *A. dorsata*, the medians of the 3 circular distributions did not significantly differ from each other (median – Start: 86.36; Middle: 91.92; End: 90.66; Fisher test statistic – Start vs Middle: 0.404, *p* = 0.525; Start vs End: 0.194, *p* = 0.659; Middle vs End: 0.017, *p* = 0.896). Finally, in *A. cerana*, the medians of the 3 circular distributions were significantly different from each other (median – Start: 79.55; Middle: 94.07; End: 104.4; Fisher test statistic – Start vs Middle: 24.725, *p* < 0.001; Start vs End: 61.418, *p* < 0.001; Middle vs End: 17.150, *p* = 0.001).

**Figure 3.**
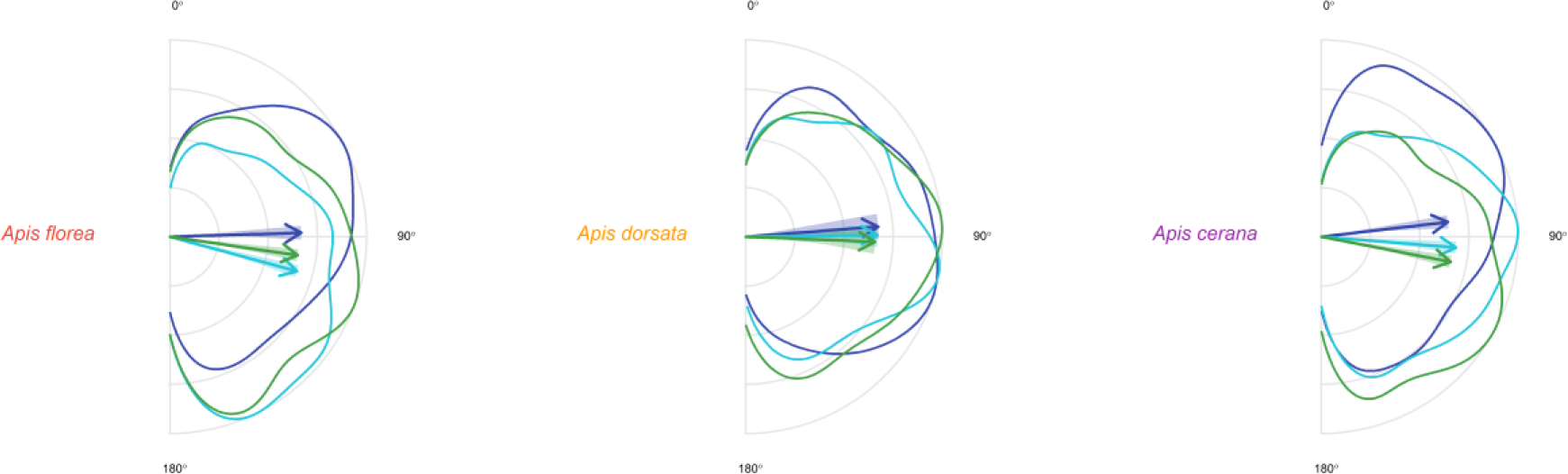
Relative circular density plots of the constrained dance follower angles for each of the different waggle run phase for *A. florea*, *A. dorsata* and *A. cerana*. The relative circular density is obtained by normalising the density in each bin of 5° to the highest density value, such that it lies between 0 and 1. The arrows in the plot represent the mean of the constrained circular distribution with the shaded region around the arrow representing the 95% confidence interval. The length of the arrow corresponds to the mean resultant length of the distribution (*ρ*). The lines representing the density plot as well as the arrows are coloured based on the waggle run phase (Start – blue, Middle – cyan, End – green).

## Discussion

The results of our study demonstrate that dance followers in three Asian honey bee species, *A. florea*, *A. dorsata* and *A. cerana* behave similarly throughout the waggle run. At the start of the run, most followers positioned themselves laterally to the dancer. Then, in the middle and the end of the run, the number of followers in the lateral and posterior positions around the dancer was similar. Further, in all three species, the mean orientation of the dance follower was close to 90° throughout the waggle run. The species differed in the number of followers per run. Dances by *A. florea* and *A. dorsata* foragers attracted larger number of followers than those by *A. cerana* foragers. The distance of the food source, and hence the duration of the waggle phase had no effect on the average number of followers present per waggle run in all 3 species.

The waggle run is hypothesized to represent a ritualization of the initiation of flight towards the food source [34–36,39]. In the open nesting and phylogenetically ancestral honey bee, *A. florea*, dances are indeed oriented in direction of the food source and there is no transposition of the direction information to a vertical plane, unlike in the giant and cavity nesting honey bees [40]. With respect to these two traits, one would predict that there should be strong differences in the dance follower behaviour between *A. florea* and the other species.

Specifically, dance followers should align themselves behind the dancers as this would allow them to most easily obtain the direction of the food source. However, in contrast to these two predictions *A. florea* dance followers neither aligned themselves behind the dancer nor showed any other major differences in their behavior in comparison with followers in the other species’. This finding supports the idea of an evolutionarily conserved mechanism of spatial information transfer in the dance behavior in all honey bees species [for *A. mellifera*, see 29].

Regarding the question of which sensory signals the dance followers use to obtain the spatial information of the dance, our results provide two arguments for the “tactile hypothesis”. The first is the higher number of followers in the lateral positions around the dancer throughout the run, which is similar to the pattern observed in *A. mellifera* [25,29,59]. The second line of evidence comes from the median body angle of the dance followers, which was close to 90° throughout the waggle run for all 3 species. Thus, followers preferred arranging themselves perpendicular to the dancer, likely using the same signals associated with this position, to obtain the spatial information in all 3 species.

Regarding the “follow hypothesis”, the pattern of the number of followers in the posterior position seen in our study provides an argument against it. The number of followers in the posterior position at the start of the run was not significantly different from the number of followers in the anterior position and was lower than the number of followers in the lateral position. Since the entire run encodes spatial information, the number of followers in the posterior position should have been high throughout the dance if following from this position was important for the information transfer. Similar to previous studies, the number of followers in the posterior position increased as the run progressed [29,60], certainly a direct consequence of the dancers forward movement during the run [29,59]. As the dancer moves forward, followers are shifted from the lateral to the posterior position. However, there was no decrease in the number of followers in the lateral position in the middle and the end of the waggle run in our observations. This suggests that either some of the followers can actively maintain their lateral positions or that vacated positions to the lateral side of the dancer are immediately occupied. Both possibilities would support the idea that the lateral position is more important than the posterior.

If the “tactile hypothesis” is correct, tactile cues associated with the lateral position around the dancer are the mechanism by which spatial information is transferred during the waggle dance. Dance followers who are laterally positioned experience a regular pattern of antennal deflections which correlate strongly with the number of abdomen waggles [29,60,61]. Since the frequency of waggling of the abdomen is likely to be similar amongst bees of the same species due to physical constraints [62], followers can use this to estimate the duration of the waggle phase. At the same time, the dance followers can obtain the orientation of the waggle run by using their own body position with respect to gravity as a reference. The Johnston’s organ may play a major role in sensing information about the direction of the waggle phase and hence the direction of the food source [63,64]. The similarity in dance follower behaviour across four species of the genus *Apis* [this study, 29] suggests that the mechanism of spatial information transfer in the waggle dance is likely through these tactile cues.

To further substantiate the “tactile hypothesis”, detailed high-speed video recordings of the antennal contacts between the dancers and the followers in all 3 species during the waggle run would be needed. In addition, the follower’s flight patterns after exiting the hive should also be observed to identify whether the information is transmitted. Even though our study provides strong support for the tactile hypothesis, we cannot completely rule out the possibility that followers can also obtain relevant information from orienting themselves behind the dancer [65]. Recent studies which have focused on the information transfer during the dance show that the transfer does not depend on the follower position around the dancer [33,65,66]. However, these studies only quantified the number of followers in the various zones around the dancer and did not compare the mean body orientation of the followers while following the dance. Combining detailed observations of the follower behaviour using a high-speed camera with tracking of their foraging trips [28] is essential to gain a better understanding of the mechanism underlying spatial information transfer in the waggle dance and confirm the tactile hypothesis.

Differences in other signals associated with the waggle dance in the various *Apis* species is linked to the modality best suited to attract followers to the dancer according to the nest environment of the species (fig 4). In our study, the two open nesting species, *A. florea* and *A. dorsata*, had higher numbers of dance followers throughout the waggle run as compared to the cavity nesting *A. cerana*. Additionally, the median body orientation did not change significantly throughout the run in the case of *A. dorsata*, while it increased from the start to the end of the run in both *A. florea* and *A. cerana*. Dancers in *A. dorsata* produce both a visual and an acoustic signal, while dancers in *A. florea* and *A. cerana* are known to use only one additional signal modality, visual and acoustic respectively [36,40,67]. Further work will be needed to tease out the exact modulatory effect of the additional signals in the waggle dance in these species.

**Figure 4.**
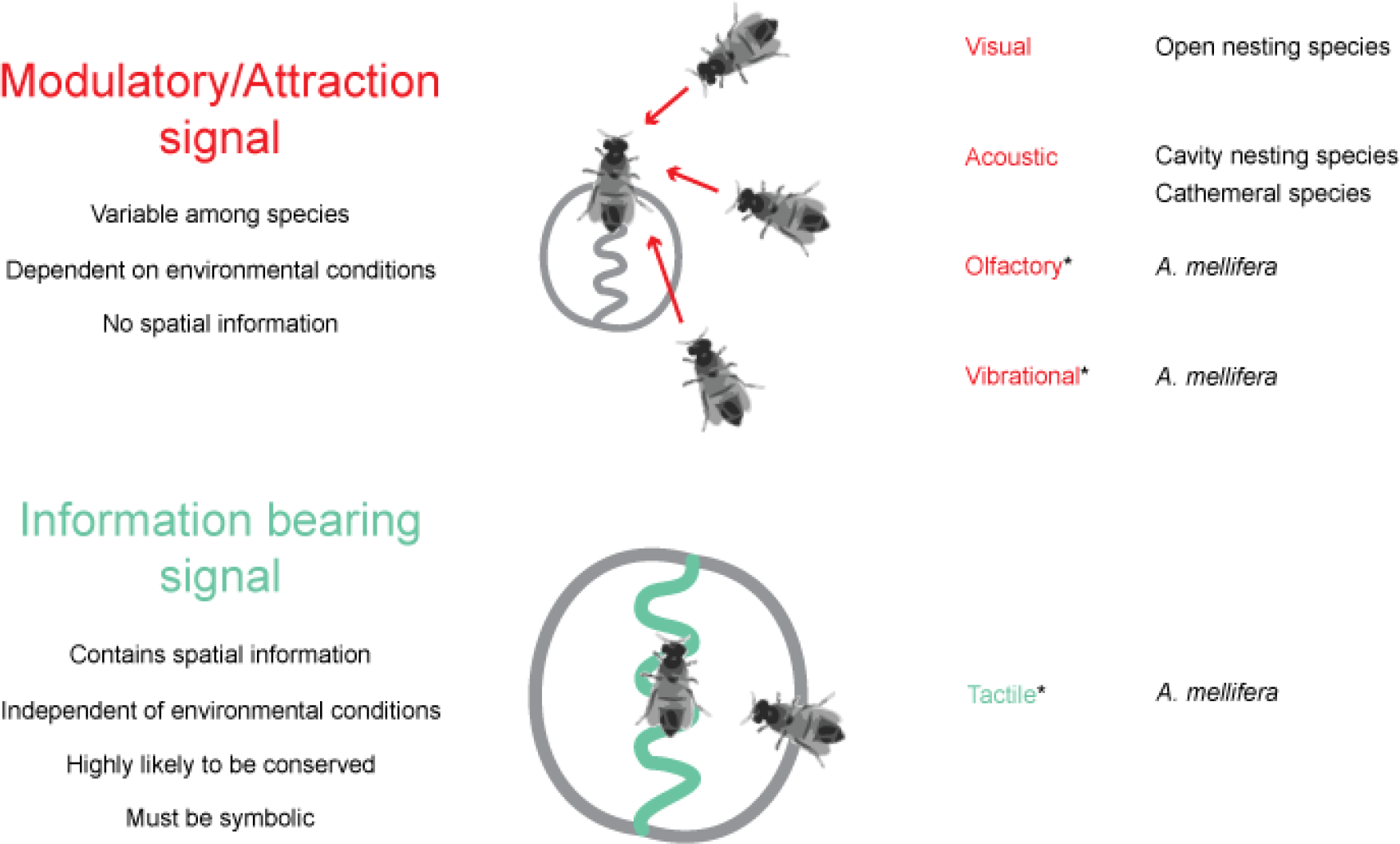
Signals produced by the dancer in different honey bee species. Signals can either attract followers to the dancer or contain information. While the former is expected to be different amongst species depending on nesting and foraging conditions, the latter should be highly conserved amongst the species. Signals with an asterisk (*) next to them have only been studied in *A. mellifera* so far. Visual and acoustic signals are only present in some species, olfactory signals cannot contain any spatial information and vibrational signals are not expected to play a role in the open nesting species as the dances often happen over other bees [34]. Tactile signals are the most likely to contain spatial information about the food source in the genus *Apis*.

In conclusion, the behavioural responses of the followers to the differing dance signals in Asian honey bees provides new insights into the evolution of this complex communication system. The difficulty in incorporating a mechanism for the transfer of spatial information into any communication system is evident from the fact that such a mechanism has evolved only rarely in animals [12]. The symbolic communication of navigational information in *Apis* likely evolved on the basis of a less complicated and non-error prone modulatory communication as seen in the closely related stingless bees and bumblebees [34–36,68]. Given the difficulty in evolving such a symbolic communication system, it can be expected that there will be very little variation in the mechanism for information transfer within a small group of closely related species. In line with this, we found that the behaviour of the dance followers, who receive the spatial information, is highly conserved across the genus. Additional signals in the waggle dance of the different species may be involved in attracting followers to the dancer (fig 4). Thus, our study highlights the usefulness of comparative studies to understand complex communication systems like the honey bee waggle dance and provides a foundation for future studies exploring the dancer and follower behaviour in the genus *Apis*.

## Data Accessibility

All the raw data obtained from these experiments is available online at Figshare (https://doi.org/10.6084/m9.figshare.11790342.v1).

## Author Contributions

E.A.G. participated in the conception and design of the study, participated in the experiments, performed the data analysis and drafted the manuscript. S. P. participated in the video and data analysis and critically revised the manuscript. N. T. participated in the experiments and the video analysis and critically revised the manuscript. A. B. participated in the conception and design of the study, coordinated the study and helped draft the manuscript.

## Acknowledgments

We are thankful to Patrick Kohl for spearheading the project on dance dialects in Asian honey bees, the videos of which we have used in this analysis and for comments on this draft of the manuscript. We are also thankful to Patrick and Benjamin Rutschmann for providing us with video recordings of dancing foragers in *A. dorsata*.

