## Supplementary Tables and Figures for "Similarities in the behaviour of dance followers among honey bee species suggest a conserved mechanism of dance communication"

E.A.G.: 0000-0002-5533-5428

S.P.: 0000-0002-6683-5946

N.T.: 0000-0003-4702-8900

A.B.: 0000-0003-0201-9656

* corresponding author:

Ebi Antony George

### Tables

#### Table S1

| Distance (m) | *Apis florea* | | | *Apis dorsata* | | | *Apis cerana* | | |
|---|---|---|---|---|---|---|---|---|---|
| | Bees | Dances | Waggle Runs | Bees | Dances | Waggle Runs | Bees | Dances | Waggle Runs |
| 100 | 5 | 9 | 65 | 1 | 1 | 47 | 4 | 8 | 94 |
| 200 | 3 | 7 | 50 | 2 | 2 | 26 | 3 | 7 | 79 |
| 300 | 7 | 9 | 65 | 2 | 2 | 28 | 5 | 7 | 84 |
| 400 | 7 | 8 | 85 | 1 | 2 | 18 | 4 | 6 | 62 |
| 500 | 5 | 8 | 65 | - | - | - | 3 | 7 | 92 |

List of bees, dances and waggle runs analysed for each of the 3 *Apis* species in this study.

#### Table S2

| Model Number | Conditional Model | | | | Zero-inflation Model | | |
|---|---|---|---|---|---|---|---|
| | Zone | Waggle Run Phase | Species | Distance | Zone | Waggle Run Phase | Species |
| 1 | + | | | | | + | |
| 2 | | + | | | + | + | + |
| 3 | | | + | | + | + | + |
| 4 | | | | + | + | + | + |
| 5 | + | + | | | + | + | + |
| 6 | + | | + | | + | + | + |
| 7 | + | | | + | + | + | |
| 8 | | + | + | | + | + | + |
| 9 | | + | | + | + | + | + |
| 10 | | | + | + | + | + | + |
| 11 | * | * | | | + | + | + |
| 12 | * | | * | | + | + | + |
| 13 | * | | | * | + | + | + |
| 14 | | * | * | | + | + | + |
| 15 | | * | | * | + | + | + |
| 16 | | | * | * | + | + | + |
| 17 | + | + | + | | + | + | + |
| 18 | * | * | + | | + | + | + |
| 19 | * | + | * | | + | + | + |
| 20 | + | * | * | | + | + | |
| 21 | + | + | + | + | + | + | |
| 22 | * | * | + | + | + | + | + |

| | | | | | | | |
|---|---|---|---|---|---|---|---|
| 23 | * | + | * | + | + | + | + |
| 24 | * | + | + | * | + | + | + |
| 25 | + | * | * | + | + | + | + |
| 26 | + | * | + | * | + | + | + |
| 27 | + | + | * | * | + | + | |

List of models used in the model selection and model averaging. All models had the number of followers as the response and were zero-inflated Poisson models with the bee ID as a random effect. The predictors present in the conditional part of each model are either the zone of the follower, the waggle run phase, the species, the distance or a combination thereof. A '+' indicates that the predictor was encoded in the model without an interaction term, whereas an '*' indicates that two predictors in the model were encoded with an interaction term between them. Only 2-way interactions were included in conditional part of the models. The predictors present in the zero-inflated part of the model were usually the zone of the follower, the waggle run phase and the species. In five models, due to overfitting, we had to reduce the number of predictors in the zero-inflated part of the model. No interactions were included in the zero-inflated part of the model.

**Table S3**

| Model Number | Degrees of Freedom | Log Likelihood | AICc | Delta AICc | Weight |
|---|---|---|---|---|---|
| *18* | *19* | *-10983.45* | *22005.0* | *0.00* | *0.662* |
| 22 | 20 | -10983.17 | 22006.5 | 1.45 | 0.321 |
| 27 | 16 | -10990.14 | 22012.3 | 7.35 | 0.017 |
| 19 | 19 | -11002.89 | 22043.9 | 38.88 | 0.000 |

List of the top four models based on their AICc values for the analysis on the number of followers. For each model, the model number (see Table S1 for predictors), the degrees of freedom, log likelihood, AICc, delta AICc and the weight is provided. Only model 18 (highlighted in italics) was present in the final shortlist of models based on a cut-off value of 0.95 of the cumulative sum of weights.

**Table S4**

| Model | Predictors | | | Effect Size | CI | *p value* |
|---|---|---|---|---|---|---|
| | Zone* | Waggle Run Phase* | Species | | | |
| Conditional Model | Anterior | Start | Apis florea | 0.2054 | 0.039 - 0.3718 | 0.0155 |
| | Lateral | Start | Apis florea | 0.4916 | 0.2371 - 0.7462 | < 0.0001 |

| | | | | | | |
|---|---|---|---|---|---|---|
| | Posterior | Start | Apis florea | 0.2625 | 0.0038 - 0.5212 | 0.2253 |
| | Anterior | Middle | Apis florea | 0.1279 | -0.1336 - 0.3893 | 0.1102 |
| | Anterior | End | Apis florea | 0.0973 | -0.1648 - 0.3594 | 0.0270 |
| | Anterior | Start | Apis dorsata | 0.2292 | -0.0124 - 0.4708 | 0.5347 |
| | Anterior | Start | Apis cerana | -0.0357 | -0.2462 - 0.1748 | < 0.0001 |
| | Lateral | Middle | Apis florea | 0.5972 | 0.1272 - 1.0673 | 0.0029 |
| | Posterior | Middle | Apis florea | 0.6309 | 0.1541 - 1.1078 | < 0.0001 |
| | Lateral | End | Apis florea | 0.6810 | 0.2106 - 1.1514 | < 0.0001 |
| | Posterior | End | Apis florea | 0.6283 | 0.1503 - 1.1064 | < 0.0001 |
| Zero-inflation Model | Anterior | Start | Apis florea | -2.7763 | -4.0726 - -1.4799 | < 0.0001 |
| | Lateral | Start | Apis florea | -16.7564 | -1908.6021 - 1875.0893 | 0.9884 |
| | Posterior | Start | Apis florea | -21.0202 | -8141.7127 - 8099.6723 | 0.9965 |
| | Anterior | Middle | Apis florea | -21.1792 | -8912.0147 - 8869.6564 | 0.9968 |
| | Anterior | End | Apis florea | -21.3387 | -9681.5457 - 9638.8683 | 0.9970 |
| | Anterior | Start | Apis dorsata | -18.2326 | -3570.3292 - 3533.8641 | 0.9932 |
| | Anterior | Start | Apis cerana | -21.7804 | -10748.4043 - 10704.8435 | 0.9972 |

List of predictors in the shortlisted model 18 for the analysis of the number of followers. For each predictor level in the final model, the effect size and confidence interval (CI) are given. In addition, the *p* value based on the comparison between the base contrast level (Anterior, Start, *Apis florea*) and the other levels obtained from the model summary is also provided [In the case of the base contrast level, the *p* value is based on whether the effect size is significantly different from zero]. The predictors marked with an * were present as an interaction term in the conditional model. In the zero-inflation part of the model, there are no interaction terms.

#### Table S5

| Species | Waggle Run Phase | $\mu$ (CI) | $\rho$ (CI) | $\beta_2$ (CI) | $\overline{\alpha_2}$ (CI) | Median |
|---|---|---|---|---|---|---|
| *Apis florea* | *Start* | *88.293 (85.2671 - 91.3188)* | *0.6676 (0.6834 - 0.6518)* | *87.466 (85.7509 - 89.1812)* | *85.6183 (83.4845 - 87.752)* | *86.69* |
| | *Middle* | *105.152 (102.449 - 107.8549)* | *0.6688 (0.6836 - 0.654)* | *96.8049 (95.282 - 98.3277)* | *84.6392 (82.6064 - 86.6719)* | *108.43* |
| | *End* | *98.2651 (95.5425 - 100.9876)* | *0.6561 (0.6703 - 0.6419)* | *93.662 (92.0888 - 95.2352)* | *86.8135 (84.9144 - 88.7126)* | *100.33* |
| *Apis dorsata* | Start | 85.6489 (80.1198 - 91.178) | 0.6755 (0.7044 - 0.6466) | 0 | 84.8189 (80.9594 - 88.6785) | 86.36 |
| | Middle | 89.3173 (84.1632 - 94.4714) | 0.6701 (0.6973 - 0.6428) | 0 | 85.1766 (81.5321 - 88.821) | 91.925 |
| | End | 92.2348 (86.8583 - 97.6112) | 0.6576 (0.6858 - 0.6293) | 0 | 86.5634 (82.8405 - 90.2862) | 90.665 |
| *Apis cerana* | *Start* | *83.3729 (80.3027 - 86.4432)* | *0.6471 (0.6626 - 0.6316)* | *85.7728 (83.9274 - 87.6182)* | *88.2383 (86.1242 - 90.3524)* | *79.555* |
| | *Middle* | *94.5799 (91.966 - 97.1939)* | *0.6858 (0.6999 - 0.6718)* | *91.4155 (90.0163 - 92.8147)* | *83.1709 (81.2688 - 85.0729)* | *94.075* |
| | *End* | *100.9708 (98.2417 - 103.6999)* | *0.6695 (0.6842 - 0.6548)* | *94.9577 (93.4502 - 96.4652)* | *84.8524 (82.8413 - 86.8636)* | *104.4* |

List of circular summary statistics for the dance follower angles in each of the 3 waggle run phase for each of the 3 species. The mean direction ($\mu$), mean resultant length ($\rho$), second central sine movement ($\beta_2$), second central cosine movement ($\overline{\alpha_2}$) along with the estimated 95% confidence intervals is provided. In addition, the median of the circular distribution is also provided. The circular distributions for each of the 3 phases for *A. dorsata* showed reflective symmetry and hence the central sine movements are set to 0. The waggle run phases within each species which significantly differed in their medians at the $p < 0.05$ level are highlighted in italics.

**Figure S1**

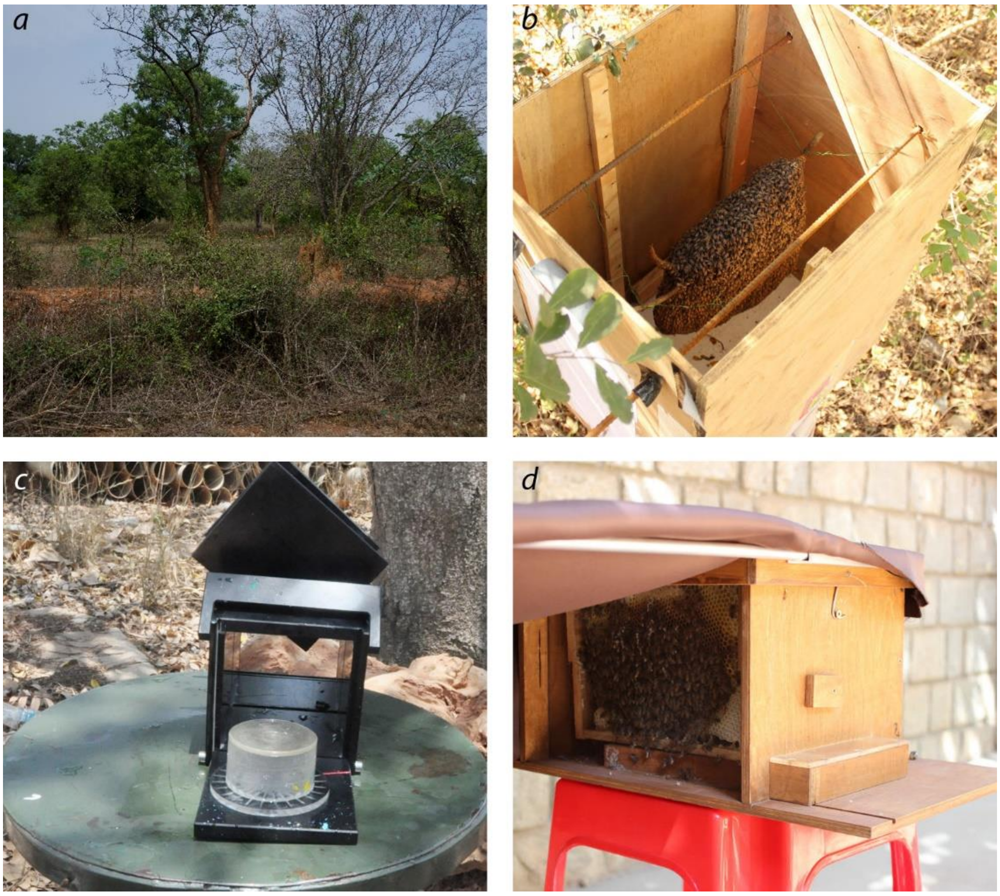


(a) A photo of the botanical garden in GKVK, Bangalore, where the experiments were performed with all the 3 species. This image is at distance of 200m from the hive in the transect used for both *A. florea* and *A. cerana*. (b) The box in which *A. florea* colonies were kept during the feeder training and dance observations. Foragers utilised the crown area of the comb as the dance floor for recruitment of the nest mates. (c) The feeder box used to train *A. dorsata* foragers. (d) The observation hive used for housing the *A. cerana* colony (picture courtesy Patrick Kohl). The observation hive had a glass wall on the side towards the entrance, which allowed us to record dances happening in the first frame (which acted as the dance floor).

Figure S2

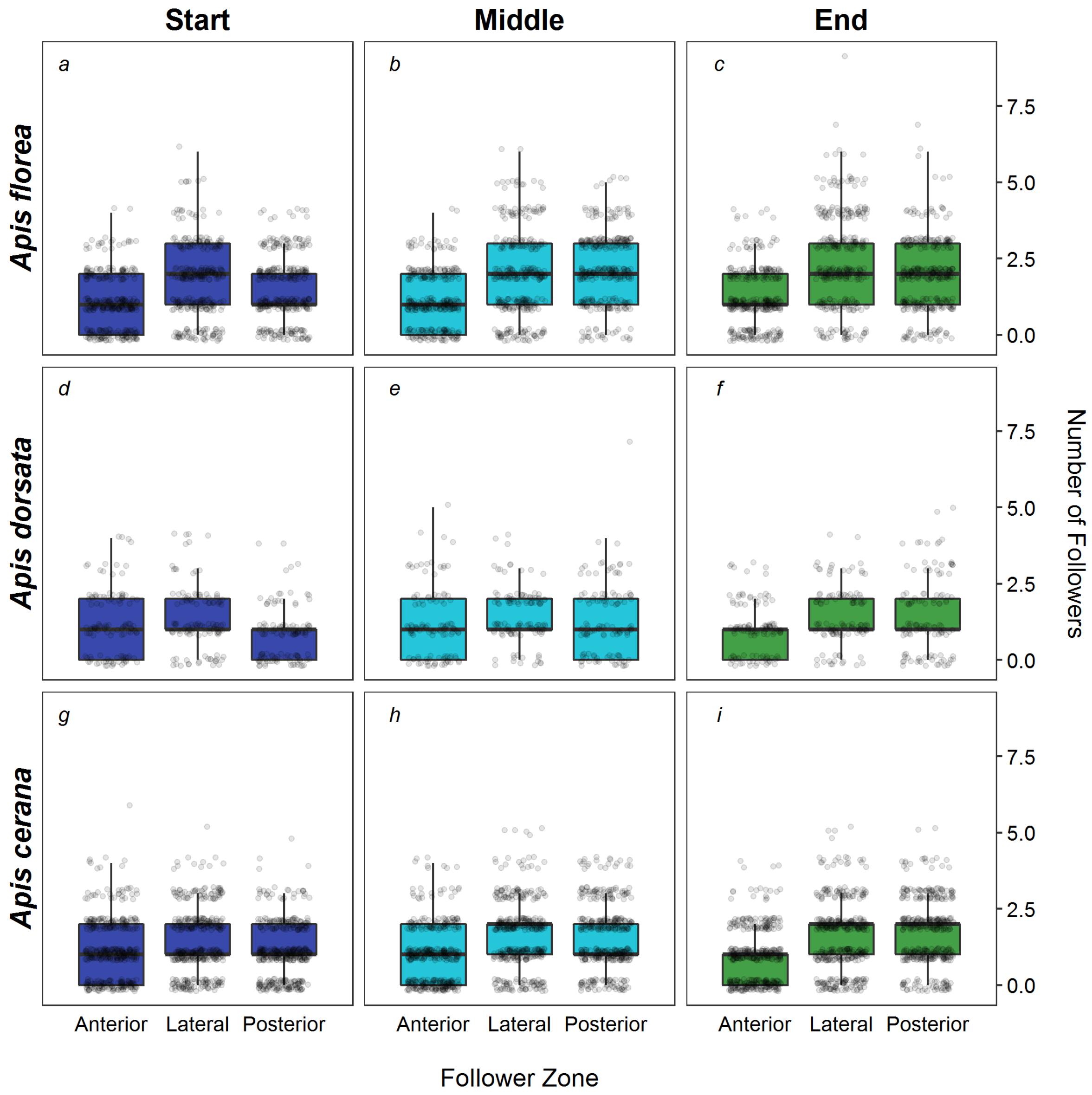


Boxplots (with the median and the quartile ranges) representing the number of followers present in each zone over the 3 phases of the waggle run for all 3 species. Each circle represents the number of followers present in one waggle run. The boxplots are filled according to the waggle run phase, with blue representing the start, cyan representing the middle and green representing the end of the waggle run.

**Figure S3**

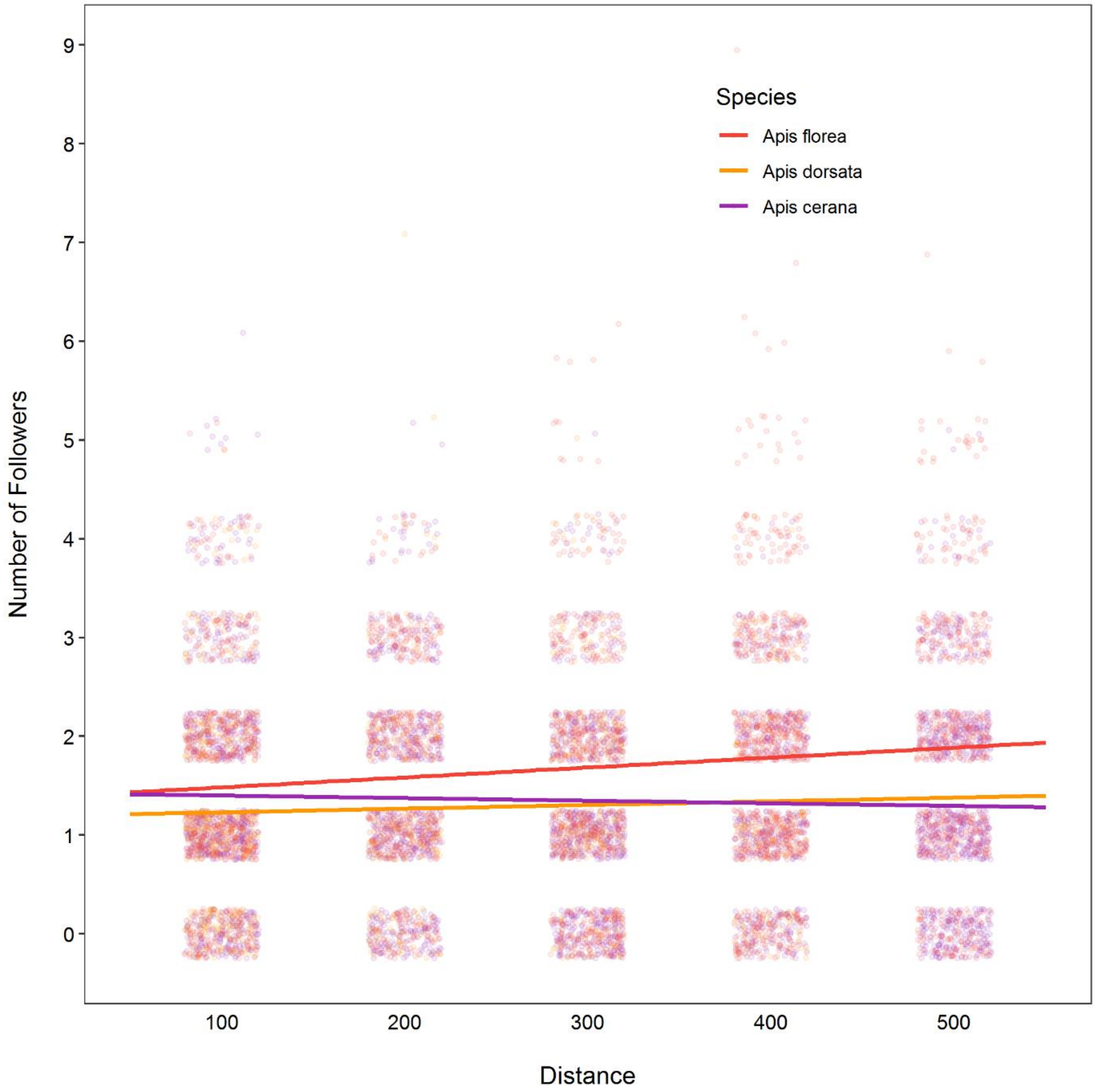


Correlation between distance and the number of followers in all 3 species. The line represents the best fitting linear correlation with each circle representing data from one run. The lines and circles are coloured according to the species, with red for *A. florea*, orange for *A. dorsata* and purple for *A. cerana*. There was no significant effect of distance on the number of followers in a waggle run in either of the 3 species, as can be seen from the horizontal lines.

**Figure S4**

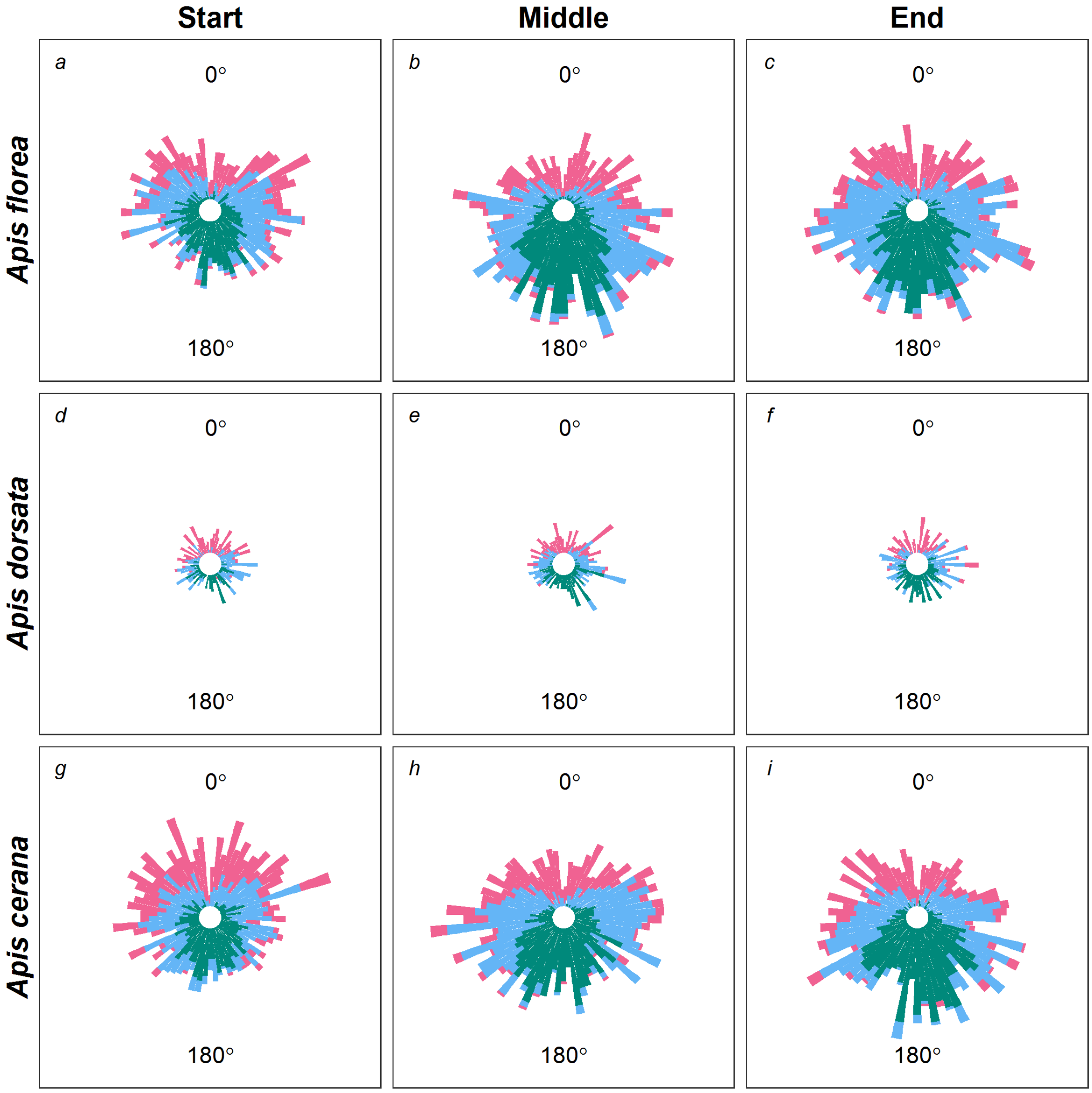


Circular histogram of the body orientation of the followers with respect to the dancer. Each bar represents the number of followers in a bin width of 5°. A follower at 0° is facing the dancer and a follower at 180° is exactly behind the dancer. The bars are coloured based on the zones occupied by the follower with pink for followers in the Anterior zone, blue for followers in the Lateral zone and green for followers in the Posterior zone.
